# Feasibility of intranasal delivery of thin-film freeze-dried monoclonal antibodies

**DOI:** 10.1101/2023.10.23.563139

**Authors:** Yu-Sheng Yu, Haiyue Xu, Khaled AboulFotouh, Gerallt Williams, Julie Suman, Sawittree Sahakijpijarn, Chris Cano, Zachary N. Warnken, Kevin C.-W. Wu, Robert O. Williams, Zhengrong Cui

## Abstract

Monoclonal antibodies (mAbs) administered intranasally as dry powders can be potentially applied for the treatment or pre-exposure prevention of viral infections in the upper respiratory tract. However, a method to transform the mAbs from liquid to dry powders suitable for intranasal administration and a device that can spray the dry powders to the desired region of the nasal cavity are needed to fully realize the potentials of the mAbs. Herein, we report that thin-film freeze-drying can be applied to prepare aerosolizable mAb dry powders and that the dry powders can be sprayed into the posterior nasal cavity using Aptar Pharma’s Unidose (UDS) Powder Nasal Spray System. AUG-3387, a human-derived mAb that neutralizes the severe acute respiratory syndrome coronavirus 2 (SARS-CoV-2), was used in the present study. First, we prepared AUG-3387 thin-film freeze-dried powders (i.e., TFF AUG-3387 powders) from liquid formulations containing different levels of mAbs. The TFF AUG-3387 powder with the highest solid content (i.e., TFF AUG-3387C powder) was then chosen for further characterization, including the evaluation of the plume geometry, spray pattern, and particle size distribution after the powder was sprayed using the UDS Powder device. Finally, the deposition patterns of the TFF AUG-3387C powder sprayed using the UDS Powder device were studied using 3D-printed nasal replica casts based on an adult model and a child model. It is concluded that it is feasible to intranasally deliver mAbs as dry powders by transforming the mAbs into dry powders using thin-film freeze-drying and then spray the powder using the UDS Powder device.

## 1. Introduction

Many viral infections, such as seasonal flu, respiratory syncytial virus (RSV) infection, the coronavirus disease 2019 (COVID-19), initiate in the upper respiratory tract, including the nasal cavity (Jain, Schweitzer and Justice, 2022; Petersen et al., 2020; Pormohammad et al., 2021; van Riel et al., 2010). Monoclonal antibodies (mAbs) that neutralize the viruses can be used to prevent or treat infections. For example, several mAb products have been authorized by regulatory agencies for COVID-19 treatment or pre-exposure prevention, including GlaxoSmithKline and Vir Biotechnology’s Sotrovimab (Heo, 2022), and AstraZeneca’s Tixagevimab-Cilgavimab (Keam, 2022). These mAbs are administered by intravenous (IV) infusion or intramuscular (IM) injection (Kelley et al., 2022). Intranasal administration of the mAbs, alone or in combination with systemic injection of them, may improve patient outcomes, and data from recent pre-clinical and clinical studies support the feasibility of delivering such mAb products intranasally to neutralize SARS-CoV-2 in a mouse model or to reduce lung inflammation and blood inflammatory biomarkers in mild to moderate COVID-19 patients (Halwe et al., 2021; Moreira et al., 2021). However, intranasal delivery of mAbs in liquid formulations has drawbacks such as the limited volume that can be administered (Li et al., 2000) and the short residence time of the liquid in the nasal cavity (Filipović-Grčić and Hafner). Intranasal administration of mAbs as dry powders may have advantages, including a prolonged residence time in the nasal cavity, benefits in storage and distribution, and the ability to modify the dissolution of the mAbs from the dry powders (Djupesland, 2013; Filipović-Grčić and Hafner; Nižić Nodilo et al., 2021).

Unfortunately, converting mAbs from liquid formulations to aerosolizable dry powders for intranasal delivery can be challenging. Conventional (shelf) freeze-drying of mAbs in the presence of various lyoprotectants such as disaccharides has proven feasible (Cleland et al., 2001; Haeuser et al., 2020), but shelf freeze-dried products are generally not suitable for aerosolization. Spray drying and spray freeze-drying can produce dry powders with desirable aerosol properties, but they are inherently associated with shear stress and high interfacial surface area during the atomization process and the heat stress unique to spray drying (Emami et al., 2018), which can potentially damage the mAbs (Pabari et al., 2011). Thin-film freeze-drying is a technology that can be applied to engineer aerosolizable dry powders while avoiding shear and heat stresses (Zhang et al., 2021). Compared to conventional shelf freeze-drying, thin-film freeze-drying provides higher freezing rate (i.e., 100-1000 K/s vs. 0.5-1 K/min) and potentially faster drying rate (Engstrom et al., 2008; Wang et al., 2023). Importantly, the highly porous, brittle matrix structures and the low bulk density of the thin-film freeze-dried powders often have desirable aerosol properties (Praphawatvet, Cui and Williams, 2022). Previously, we have demonstrated that it is feasible to apply thin-film freeze-drying to prepare dry powders of mAbs for pulmonary delivery by oral inhalation (Emig et al., 2021; Hufnagel et al., 2022; Sahakijpijarn et al., 2020). For example, AUG-3387, a human-derived mAb that neutralizes SARS-CoV-2, was successfully thin-film freeze-dried into aerosolizable dry powders for pulmonary delivery (Emig et al., 2021). Herein, we tested the feasibility of using the thin-film freeze-drying technology to prepare dry powders of mAbs suitable for intranasal delivery. Again, AUG- 3387 was used, though in modified formulations. The resultant thin-film freeze-dried AUG-3387 mAb powders (TFF AUG-3387 powders) were characterized, and their deposition patterns in 3D-printed nasal replica casts were studied using Aptar Pharma’s UDS Powder device.

## 2. Materials and Methods

### 2.1. Materials

AUG-3387 mAb was from TFF Pharmaceuticals, Inc. (Austin, TX). L-histidine and fluorescein isothiocyanate isomer I (FITC) were from Acros Organics (Geel, Belgium). D-mannitol, Bradford reagent, and Tween 20 were from Sigma-Aldrich (St. Louis, MO). Leucine was from Spectrum Chemical Mfg. Corp. (New Brunswick, NJ). Artificial nasal mucus was from Biochemazone (Leduc, Canada).

### 2.2. Preparation of AUG-3387 dry powders

AUG-3387 was dialyzed overnight in a histidine buffer (20 mM, pH 6) containing 0.02% of Tween 20. The composition of the histidine buffer was based on Haeuser et al. (2020). Then, histidine buffer containing mannitol and leucine was added to the mAb solution according to the ratios in Table 1. Histidine buffer was used to adjust the final volume. The liquid mAb formulations were then converted to powders by thin-film freeze-drying. Briefly, the liquid formulations were dropped onto the surface of a rotating cryogenically cooled stainless steel drum with a BD 1 mL syringe equipped with a 21G needle. The temperature of the drum was controlled at -70 to -100°C. The frozen films were collected into a container containing liquid nitrogen and transferred to 5 mL glass vials. The vials were semi-stoppered, and the formulations were dried using a SP Scientific Virtis Advantage Pro lyophilizer (Warminster, PA). The lyophilization process consisted of a 20 h primary drying step at -40°C, a ramping step from -40°C to 25°C over 20 h, and a 20 h secondary drying step at 25°C. The chamber pressure of the lyophilizer was maintained at 80 mTorr. After the lyophilization process, the vials were back-filled with nitrogen, stoppered, crimped, and then stored at room temperature.

**Table 1.** AUG-3387 mAb formulations with different solid contents. The final volume was adjusted to with a histidine buffer (20 mM, pH 6) containing 0.02% of Tween 20.

|  | AUG-3387A | AUG-3387B | AUG-3387C |
| --- | --- | --- | --- |
| AUG-3387 (g/L) | 1.49 | 2.98 | 4.47 |
| Mannitol (g/L) | 8.03 | 16.06 | 24.09 |
| Leucine (g/L) | 0.423 | 0.846 | 1.269 |
| Final volume (μL) | 500 | 250 | 167 |

### 2.3. Size exclusion chromatography analysis

An Agilent 1260 Infinity II liquid chromatography (LC) system (Santa Clara, CA) equipped with an AdvanceBio SEC 300 Å 2.7 µm 7.8 × 300 mm column was used to evaluate mAb monomer and high molecular weight (HMW) species. The mAb samples were diluted to 0.298 mg/mL, filtered through a 2.0 µm glass microfiber syringe filter and a 0.45 µm polyethersulfone (PES) syringe filter, and stored in a refrigerator. The chromatographic conditions were: sample volume, 5 µL; mobile phase, 0.15 M phosphate buffer (pH 7); run time, 40 min; flow rate, 0.5 mL/min; and detection wavelength, 220 nm. The peak area of the mAb on the chromatogram was determined using Agilent OpenLab Software, and the numerical data of the chromatogram was exported with an UniChrom software (https://www.unichrom.com).

### 2.4. Micro-flow imaging (MFI) analysis

MFI was applied to analyze proteinaceous particles in the mAb formulations. The TFF mAb formulations were first reconstituted to the original volume with water. Histidine buffer was then used to dilute the samples to reach a final mAb concentration of 0.3104 mg/mL. The samples were analyzed using an MFI 5100 micro-flow imaging system (ProteinSimple, San Jose, CA) with an auto-sampler. The analyzed sample volume was 0.35 mL. The spread sheet containing the equivalent circular diameter (ECD), area, perimeter, circularity, aspect ratio, and intensity standard deviation profiles of each detected particles were exported with the ProteinSimple MVSS software. To exclude the contaminant particles such as silicone oil droplets and air bubbles, a filter was applied to exclude the particles with the aspect ratio ≥ 0.8 and intensity standard deviation > 100 according to the literature (Guo et al., 2022). Also, the particles presence on the edge of the images were removed, while the “remove slow and stuck particles” feature was not applied. The particle counts were converted to mass of proteinaceous particles following the ellipsoid-volume (E-V) method by Kalonia et al. (2015). Briefly, the mass of an individual particle *k* (*mk*) was first determined using following equation (1):

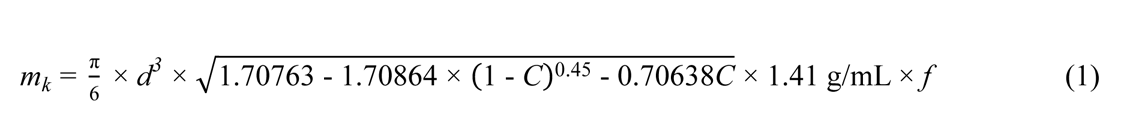

Where *d* is the ECD of the particle, *C* is the circularity of the particle, and *f* is the protein fraction of the particle. A value of 0.2 was chosen for *f* by assuming proteinaceous particles contain 20% protein and 80% solvent.

The total mass of proteinaceous particles (*M*) was then calculated following equation (2):

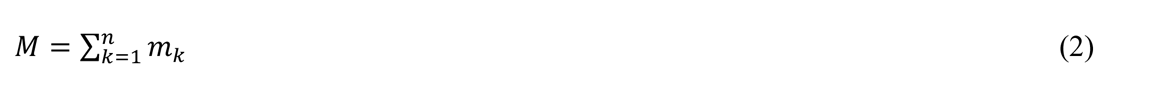

Where *n* is the number of particles.

Finally, the mass percentage of the particles was calculated using equation (3), in which we divided the *M* with the theoretical concentration of mAb in the solution (i.e., 0.3104 mg/mL) multiplied by the volume being analyzed (*Va*).

Mass percentage of the particles (%) =

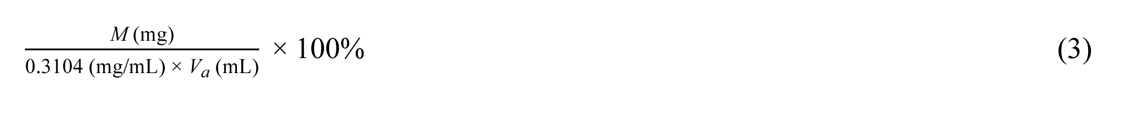

### 2.5. Measurement of the water content

The water contents in the TFF AUG-3387 powders were determined by Karl Fischer titration using a Mettler Toledo C20 coulometer (Columbus, OH). Briefly, 1 mL of the HYDRANAL™-Coulomat AG solution (Honeywell, Charlotte, NC) was drawn from the buffer tank and injected into the vial containing the TFF mAb powder. The TFF AUG- 3387C powder was dissolved with the Coulomat AG solution, and 0.5 mL of the mixture was injected back into the buffer tank. The water content (% *w/w*) of the sample was then calculated using equation (4):

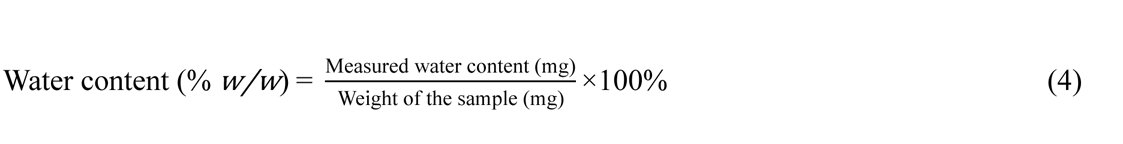

### 2.6. Measurement of the specific surface area

The specific surface area of the TFF mAb powder was measured using a Brunauer-Emmett-Teller (BET) surface area analysis. The sample was loaded into the sample cells and outgassed under helium flow for 1 day. The nitrogen absorption was then performed with p/p0 ranging from 0.1 to 0.3 with an Anton Paar Quantachrome AutoFlow BET+ instrument (Graz, Austria). The specific surface area of the TFF mAb powder was then calculated with the BET method.

### 2.7. X-ray diffraction (XRD)

The TFF AUG-3387C powder was mounted with mineral oil and loaded onto a sample loop. The 2D XRD pattern of the sample was measured with a Rigaku Spider instrument (Tokyo, Japan). The background of the sample’s 2D diffraction pattern was subtracted using the diffraction pattern of mineral oil. The 2D diffraction pattern was then transformed into 1D pattern and the 1D pattern was smoothed with the Rigaku 2D Data Processing (2DP) software. The crystallographic information files (CIF) of the mannitol (Fronczek, Kamel and Slattery, 2003), leucine (Coll et al., 1986), and histidine (Madden et al., 1972; Madden, McGandy and Seeman, 1972) were obtained from the Cambridge Crystallographic Data Centre (CCDC) (https://www.ccdc.cam.ac.uk), and the simulated XRD patterns were generated with the CCDC mercury software.

### 2.8. Plume geometry and spray pattern

TFF AUG-3387C powder (4.73 ± 0.59 mg, n = 6) was loaded to the Aptar UDS Powder device and sprayed upward manually. A custom built system containing a laser light sheet and a high-speed digital camera were used to visualize both plume geometry and spray pattern (Warnken et al., 2018). When visualizing the plume geometry, the laser light sheet was aligned with the axis of the spray, and the plumes were captured at a frame rate of 500 frames per second (FPS). The stacking images over 11 frames of the sprays were created, and the plume angles were measured using the FIJI software. When visualizing the spray patterns, the laser light sheet was perpendicular to the axis of the spray and the distance between the laser light sheet and the UDS Powder device nozzle was 6 cm. The spray patterns were then captured at a frame rate of 500 FPS and an angle of about 45°. The stacking images over 11 frames of the sprays were created by using Fiji software. The perspective correction of the images was performed using MATLAB 2022a software. Finally, Fiji software was used to determine the ovality and the area of the spray patterns. The experiment was repeated three times for both plume geometry and spray pattern.

### 2.9. Scanning electron microscopy

The scanning electron microscopic (SEM) images of the TFF AUG-3387C powder were taken with a Hitachi S-5500 field emission SEM instrument (Tokyo, Japan). The bulk TFF powder was fixed on the sample stub with a carbon tape. The sample was coated in a sputter coated equipped with an Au/Pd (60:40) target from Electron Microscopy Sciences (Hatfield, PA). The SEM images were then acquired with an acceleration voltage of 30 kV. SEM was also used to examine TFF AUG-3387C powder after it was sprayed with the UDS Powder device. The TFF mAb powder was loaded into the UDS Powder device and sprayed onto a sample stub covered with a non-porous carbon tape at a distance of 6 cm. The sample was sputtered with Au/Pd and then observed with an acceleration voltage of 30 kV.

### 2.10. Measurement of the particle size

A Malvern Spraytec laser diffraction instrument (Malvern, United Kingdom) was utilized to measure the size of the TFF mAb powder after it was sprayed using the UDS Powder device. The particle size was measured using either a ‘wet’ method or a ‘dry’ method. The procedure for the ‘wet’ method is as follows, while the details of the ‘dry’ method are in the Supplementary Material. TFF AUG-3387C powder (5.66 ± 0.29 mg, n = 2) was loaded into a UDS Powder device and sprayed into a glass test tube. The sprayed powder was dispersed with hexane, and the particle size distribution was measured using the Spraytec Wet Dispersion Accessory. During the measurement, the dispersion was driven by a mechanical stirrer with a rate of 3000 rpm and circulated through the sample cell. The sample was measured at a rate of 1 Hz, and the particle size distribution (concentration weighed) was calculated by averaging the scatter data over 10 s.

### 2.11. Intranasal deposition patterns of the TFF mAb powder

FITC-labeled AUG-3387 mAb was used to prepare the TFF AUG-3387C powder for the intranasal deposition studies. AUG-3387 was labeled with FITC following the instruction of a Sigma-Aldrich FluoroTag conjugation kit, and the product was purified by ultrafiltration. The TFF AUG-3387C powder (5.29 ± 0.75 mg, n = 12) was loaded into the UDS Powder device and sprayed into the nasal replica casts 3D-printed based on the CT-scan images of the nasal cavities of a 48-year-old male (adult) or a 7-year-old female (child) (Warnken et al., 2018). Before spraying the powder, the nasal replica casts were coated with artificial nasal mucus. A Cytiva 2.0 µm glass microfiber syringe filter (Marlborough, MA) was connected after the nasopharynx part. The powder was sprayed into the left nostrils of the replica casts. The coronal angel was fixed at 0°, and the sagittal angle was fixed at 45° based on our previous study (Yu et al., 2023). The insertion depth for was 0.5 cm for the adult nasal cast and 0.4 cm for the child nasal cast. To study the effect of the flow rate on the deposition of the TFF mAb powder, a flow rate of 0 or 10 liters per minute (LPM) was applied for the adult nasal cast, 0 or 5 LPM for the child nasal cast. After the powder was sprayed, the replica casts were disassembled into anterior, upper, middle, and lower turbinate, nasopharynx, and filter parts. Each part was rinsed with 5 mL of water, and the fluorescence intensity of the FITC-labeled AUG-3387 (λEX = 485 nm, λEM = 528 nm) was quantified using a BioTek Synergy Microplate Reader (Winooski, VT). A calibration curve was used to convert the fluorescence intensity into the amount of powder collected.

### 2.12. Statistical analyses

All the data are presented as mean ± standard deviation (SD). One-way analysis of variance (ANOVA) followed by Tukey’s post hoc test was performed using Microsoft Office Excel with the Real Statistics Resource Pack (Release 8.5) (https://real-statistics.com).

## 3. Results and discussion

### 3.1. Preparation and characterization of TFF mAb powders

Previously we have thin-film freeze-dried AUG-3387 mAb with mannitol and leucine (95:5) as the excipients into dry powders with desired aerosol properties for pulmonary delivery into the lungs, while preserving its integrity and activity (Emig et al., 2021). In this work, we modified the composition slightly by changing the buffer from the phosphate-buffered saline (PBS) to a histidine buffer and including a small amount of Tween 20 (Table 1). Three liquid formulations, AUG-3387A, AUG-3387B, and AUG-3387C, were prepared with final mAb concentrations at 1.49 g/L, 2.98 g/L, or 4.47 g/L, respectively, while the concentration of the buffer and Tween 20 remained the same. The mAb liquid formulations were then subjected to thin-film freeze-drying to produce three different dry powders. The water contents in the TFF mAb powders ranged from 1.68 to 2.32% *w/w*. Aggregation of protein-based therapeutic agents such as mAbs after they are subjected to freezing and/or drying stresses is a key concern. Therefore, we evaluated the formation of high molecular weight (HMW) species after the AUG-3387 mAb formulations were subjected to thin-film freeze-drying using SEC. HMW species were not detectable in the AUG-3387 liquid formulation (**Fig. 1**) and subjecting the mAb to thin-film freeze-drying did not result in the formation of HMW species in any of the AUG-3387 formulations (**Fig. 1**), indicating that the TFF process did not cause the formation of significant HMW species in any of the AUG-3387 formulations.

**Fig. 1.**
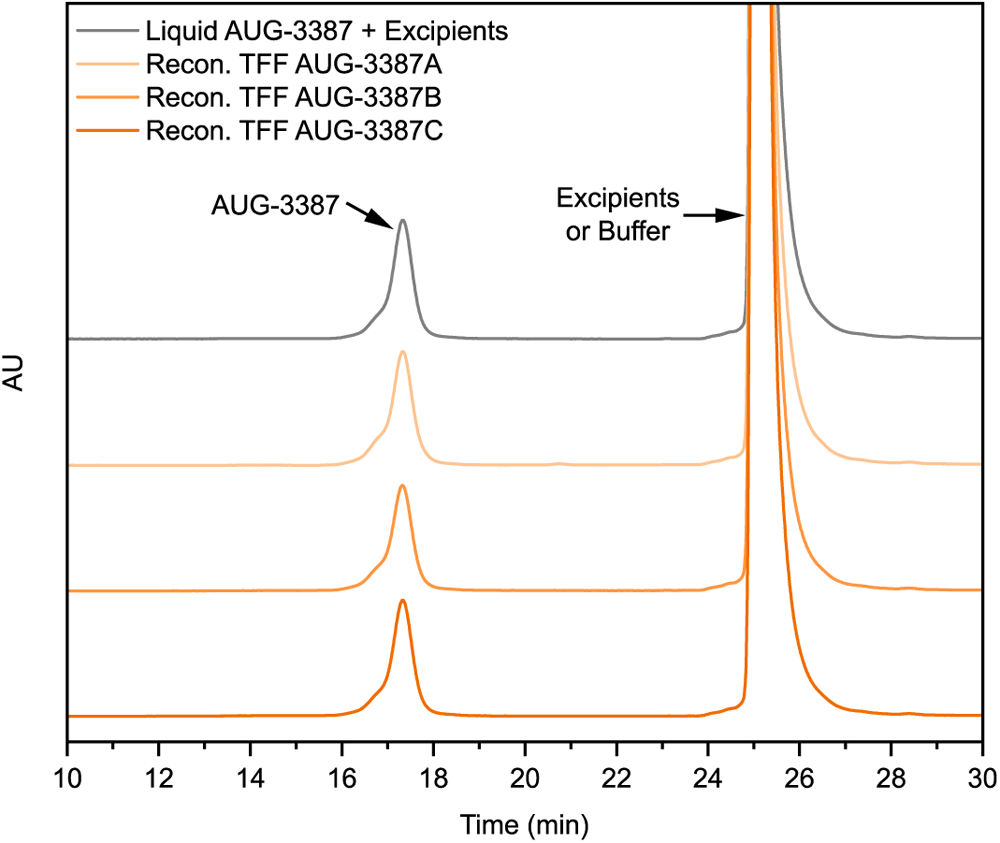
SEC chromatograms of the AUG-3387 mAb in liquid formulation or reconstituted from TFF mAb powders. The experiment was repeated three time with similar results.

Recently, MFI has emerged as an attractive method to quantify the proteinaceous particles in protein formulations (Sharma et al., 2010a). By combining a digital microscope with a microfluidic system, the images of each particle in a protein formulation can be captured, and the information of each particle such as ECD, circularity, and aspect ratio can be calculated with an image analysis software (Sharma et al., 2010b). Particles of larger than 100 μm are considered visible, and those that are smaller than 100 μm are considered subvisible. In products such as injectable protein products, there are recommended limits for the number of particles larger than 10 μm and larger than 25 μm (Glücklich et al., 2020; USP-NF, 2023), though such limits are unknown for products intended for intranasal administration. For all the powders, most of the subvisible particles were in the ranges of 2 ≤ x < 5 µm and 5 ≤ x < 10 µm. Subjecting the mAb formulations to the TFF process caused an increase in the concentration of subvisible particles, but the concentrations of subvisible particles in all three TFF AUG-3387 powders were not different (data not shown). At least two methods have previously been applied to calculate the mass of the proteinaceous particles using particle count data from MFI. In the method by Barnard et al. (2011), it is assumed that the proteinaceous particles are spherical, the volume fraction of the proteins in the particles is 75%, and the density of the protein is 1.43 g/mL. However, this method often overestimates the mass of protein aggregates, and the sum of the calculated mass is usually higher than the experimentally determined value (Kalonia et al., 2015). In the E-V method reported by Kalonia et al. (2015), the 2D images of the proteinaceous particles are converted into the prolate ellipsoids of revolution, and the masses of the particles are then calculated by assuming the volume fraction of the proteins in the particles is 20% and the density of the protein is 1.41 g/mL. We estimated the mass of proteinaceous particles in each of the TFF mAb powders following Kalonia et al. (2015). As shown in **Fig. 2**, the mass percentage of the particles in all three TFF AUG-3387 mAb powders was lower than 2%, and there was not a significant difference among those three TFF mAb powders in particle mass.

**Fig. 2.**
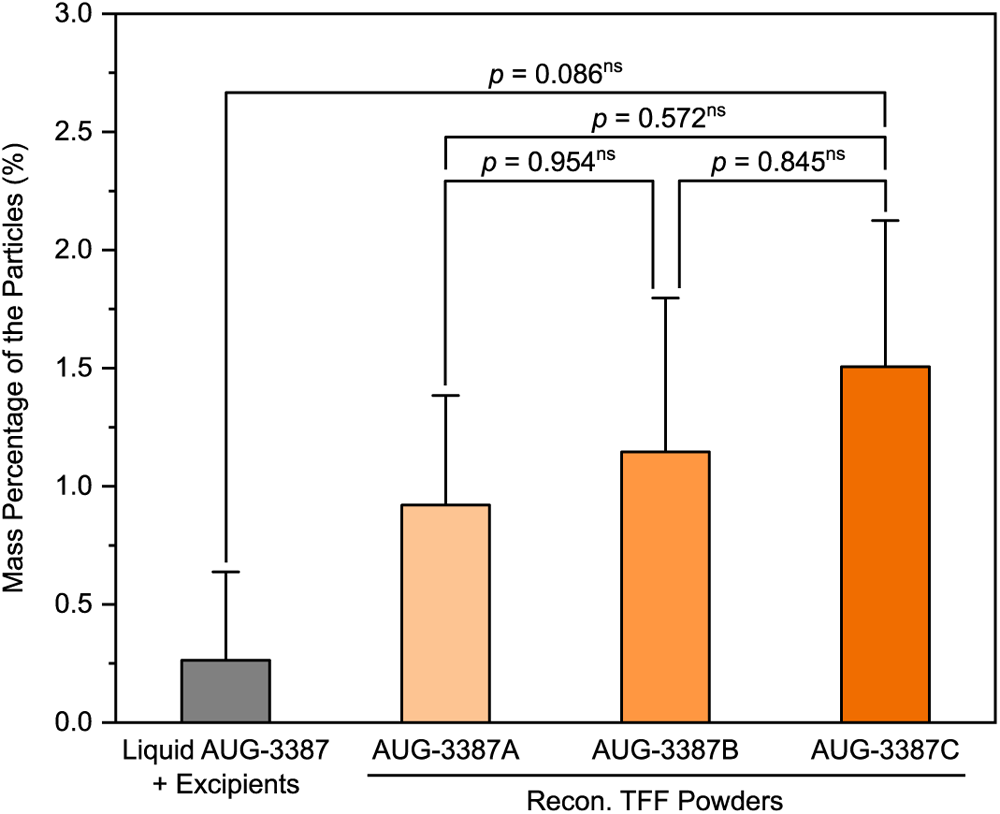
Mass percentages of particles in different TFF mAb formulations as calculated using the E-V method. Data are mean ± SD (n = 3).

According to the Karl Fischer titration, the water contents of the TFF AUG-3387A, AUG-3387B, AUG-3387C powders were 1.6 ± 0.3% *w/w*, 2.0 ± 0.5% *w/w*, and 2.3 ± 0.7% *w/w*, respectively. Since all three powders were similar in their moisture contents and the concentrations of HMW species and the proteinaceous particle masses were not different among them upon reconstitution, we decided to use AUG-3387C for additional studies. The TFF AUG-3387C powder also had the highest mAb content, allowing the filling of more mAbs in a device for intranasal spraying.

The average BET specific surface area of the TFF AUG-3387C powder was 24.934 ± 3.354 m^2^/g, indicating that it was porous. **Fig. 3** shows the X-ray diffractogram of the TFF AUG-3387C powder, in which most of the diffraction peaks match the simulated peaks of the crystalline mannitol (δ form). The peaks of crystalline leucine are not easily identifiable, although freeze-dried leucine is usually crystalline, likely because leucine accounted for only 5% of the excipients. The histidine from the histidine buffer was also amorphous in the TFF AUG-3387C powder. The diffraction patterns of two possible form of histidine crystals were simulated to confirm the absence of crystalline histidine (**Fig. 3**). It is generally believed that crystals could damage mAbs during freezing. It is unclear why the mannitol crystals did not cause more aggregation of the AUG-3387 mAbs than observed, although the “particle isolation hypothesis” theorizes that mannitol may offer partial protection against particle aggregation by separating particles from one another within the unfrozen fraction during freezing (Allison, Molina and Anchordoquy, 2000). Of course, histidine in the formulation is a known cryoprotectant (Mohammed, Coombes and Perrie, 2007).

**Fig. 3.**
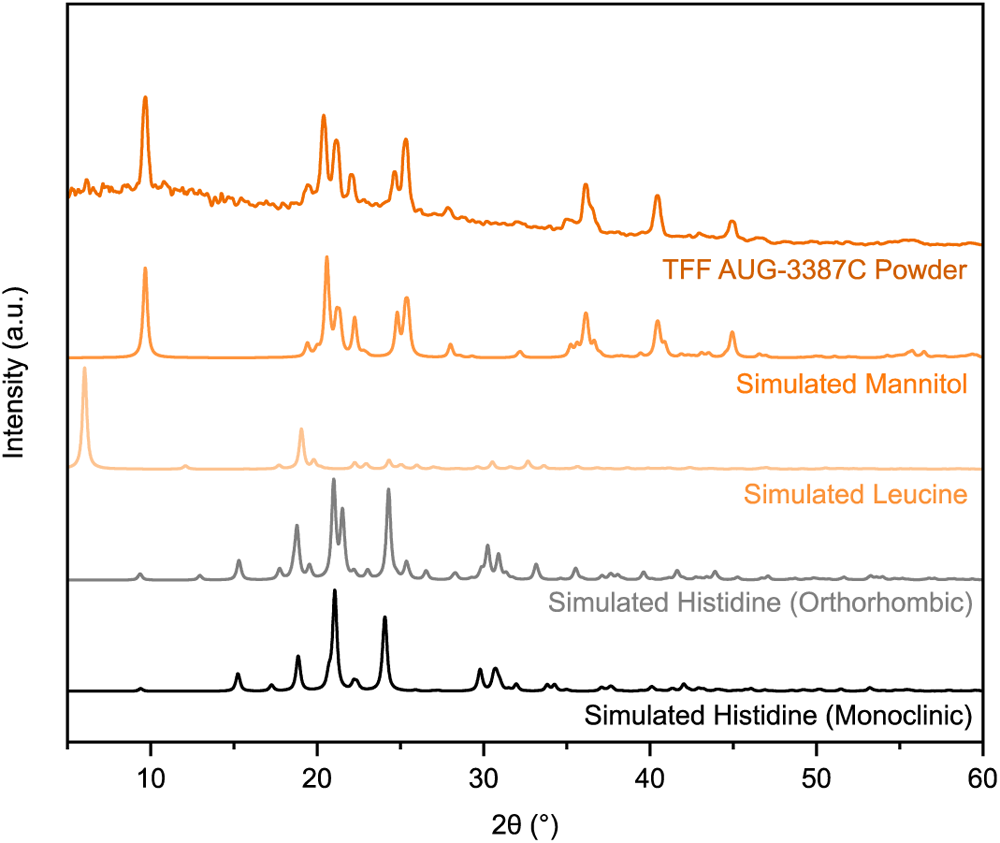
The XRD diffractograms of the TFF AUG-3387C powder and the simulated patterns of mannitol (δ form) (CCDC: 224660), leucine (CCDC: 1206031), and L-histidine (Orthorhombic, CCDC: 1206541; Monoclinic, CCDC: 1206538).

To study the plume geometry and spray pattern of the TFF AUG-3387C mAb powder, about 5 mg of the TFF mAb powder were loaded into an Aptar’s UDS Powder device and sprayed. The plume geometry and the spray pattern are shown in **Fig. 4**. For the plume geometry, the average plume angle was 16.13 ± 2.08° (n = 3). For the spray pattern, the average area was 423.73 ± 28.88 mm^2^, and the ovality was 1.20 ± 0.06 (n = 3).

**Fig. 4.**
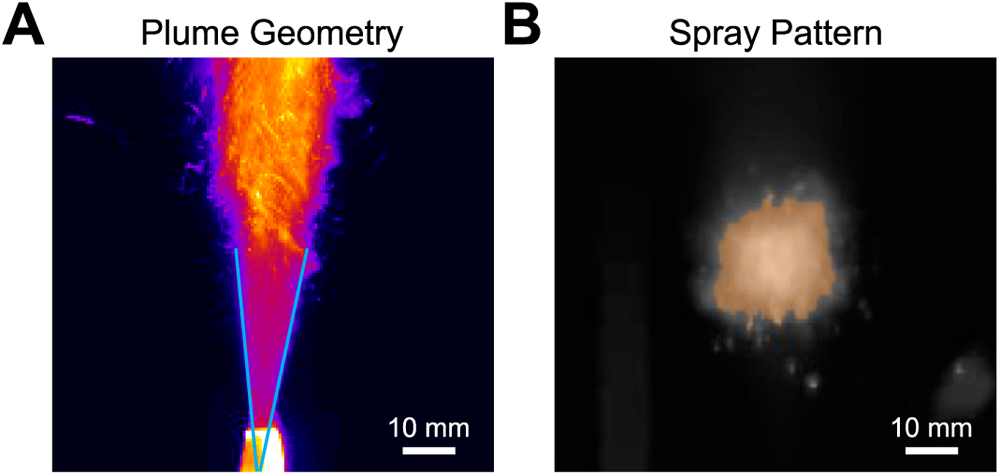
Representative plume geometry (A) and the spray pattern (B) images of the TFF AUG-3387C powder after being sprayed with the Aptar’s UDS Powder device.

The morphology of the TFF AUG-3387C powder, before and after being sprayed with the Aptar’s UDS Powder device, was examined using SEM. The TFF mAb powder was porous (**Fig. 5A**), which explains its relatively high specific surface area. After the powder was sprayed with the UDS Powder device, irregular shape particulates appeared (**Fig. 5B**), likely because the dried thin films were sheared into smaller irregular pieces during the actuation/spraying process.

**Fig. 5.**
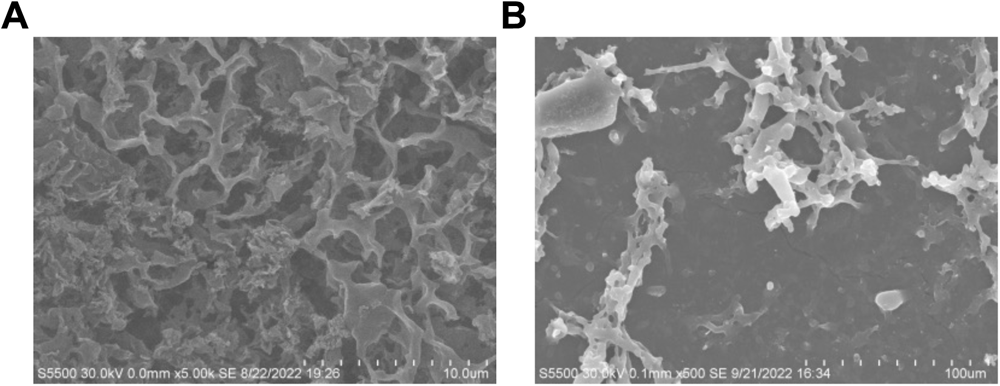
Representative SEM images of the TFF AUG-3387C powder (A) before and (B) after it was sprayed using an Aptar’s UDS Powder device.

The particle size and size distribution of TFF AUG-3387C powder after being sprayed using the UDS Powder device and then dispersed in hexane were evaluated using a Spraytec laser diffraction system and shown in **Fig. 6**. Per U.S. FDA Guidance for Industry, the droplet/particle size distribution of nasal spray and spray drug products is considered one of the most critical attributes (US FDA, 2002). When measuring droplet/particle size distribution of sprayed liquid using laser diffraction with Spraytec, the particle size within the stable phase, instead of the formation and dissipation phases is considered most important in defining the bioavailability of the intranasally delivered products (Malvern Panalytical, 2005). However, for nasal powders such as the TFF mAb powder sprayed using a device such as Aptar’s UDS Powder, a stable phase is difficult to define and the transmission is lower than 5% (data not shown), spraying the powders in to a container and dispersing them using a liquid, in which they do not dissolve, and then measuring the distribution of the dispersed particles may represent an alternative method to accurately estimate the particle size and particle size distribution of the sprayed powders.

**Fig. 6.**
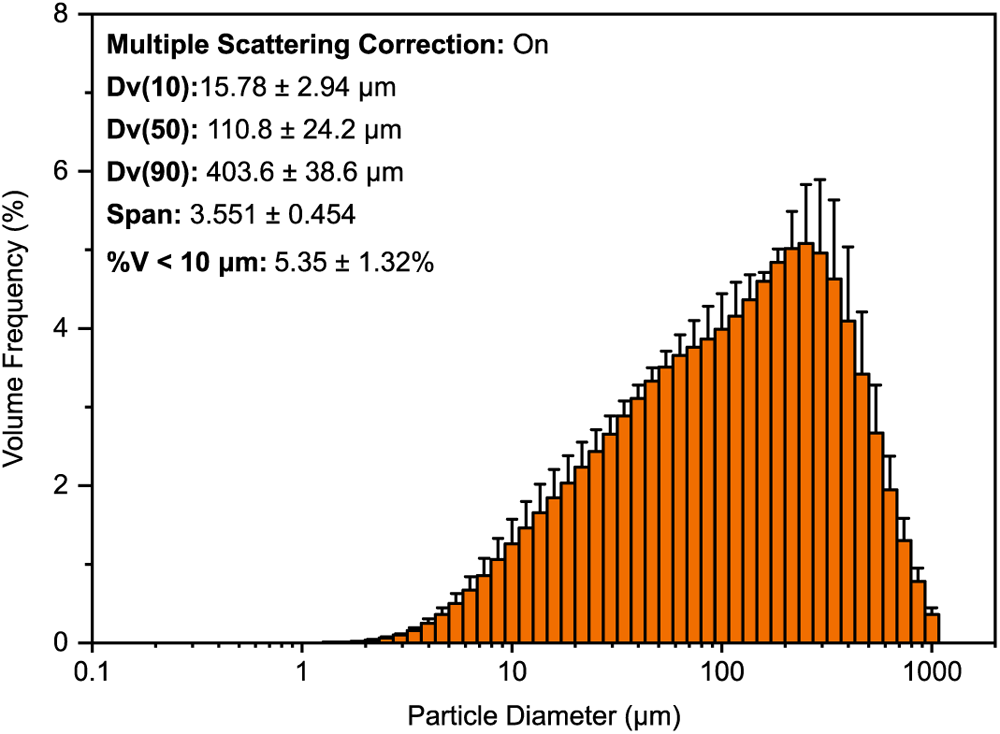
Particle size distribution in the sprayed TFF AUG-3387C mAb powder dispersed in hexane (n = 2). The particle size distribution of the sprayed powder was calculated by integrating 10 s of the signal.

### 3.2. Deposition patterns of TFF AUG-3387C powder in nasal replica casts

The deposition patterns of the TFF AUG-3387C powder in nasal replica casts based on the CT scans of an adult and a child were studied using FITC-labeled AUG-3387. **Fig. 7A** shows the deposition pattern of the TFF AUG-3387C powder in the nasal replica cast of a 48-year-old male. Most of the powder was deposited in the middle and lower turbinates and the nasopharynx region. Also, when a flow rate of 10 LPM was applied, the deposition in the nasopharynx increased, but the deposition in the middle and lower turbinates and the nasopharynx region together (i.e., RDP) did not improve (from 83.0% to 79.0%) (**Fig. 7B**). With or without applying a flow rate, the recovery percentage of the powder from the nasal replica cast was close to 100% (**Fig. 7C**).

**Fig. 7.**
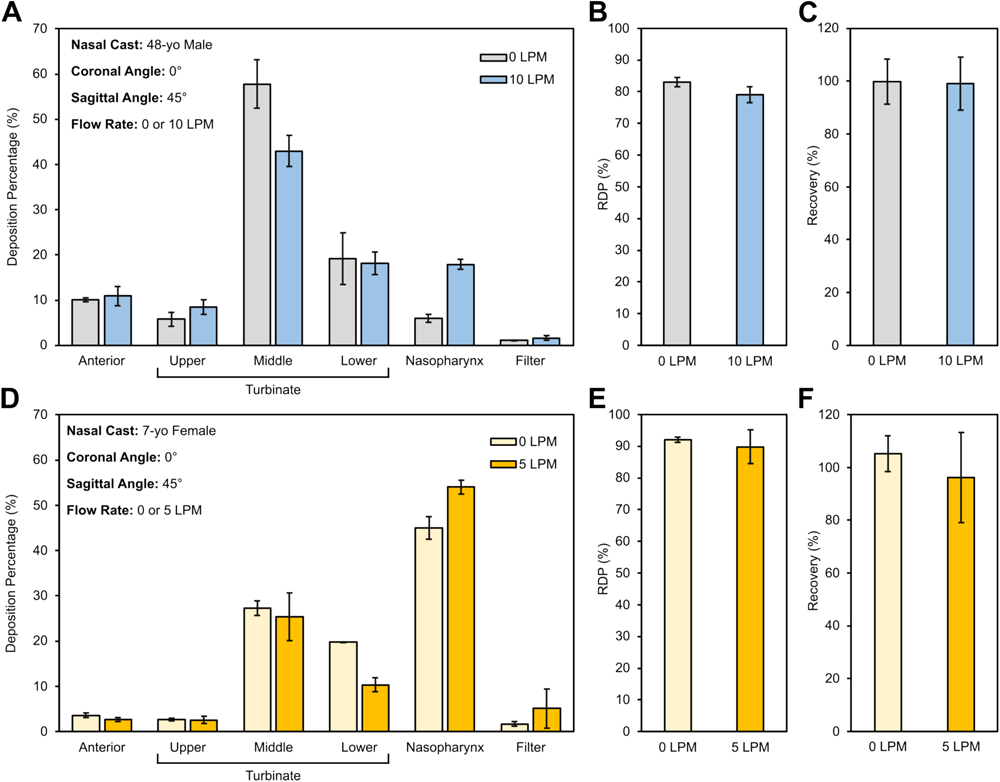
Deposition patterns of TFF AUG-3387C powder in nasal replica casts. (A) Deposition pattern of TFF AUG-3387C powder in the nasal replica cast based on the CT scan of a 48-year-old male. (B) The RDP of the powder in the 48-year-old male nasal replica cast. (C) The recovery percentage of the powder from the 48-year-old male nasal replica cast. (D) Deposition pattern of TFF AUG-3387C powder in the nasal replica cast based on the CT scan of a 7-year-old female. (E) The RDP of the powder in the 7-year-old female nasal replica cast. (F) The recovery percentage of the powder from the 7-year-old female nasal replica cast. Data are mean ± SD (n = 3).

**Fig. 7D** shows the deposition pattern of the TFF AUG-3387C powder in the nasal replica cast of a 7-year-old female. Again, most of the powder was deposited in the middle and lower turbinates and the nasopharynx region, although the deposition in the nasopharynx region was the highest. Applying a flow rate of 5 LMP led to a slight reduction in the RDP (92.1% to 89.7%) (**Fig. 7E**), because more powders went through the nasal cast and were caught on the filter. Finally, the recovery percentages of the powders in the child nasal replica cast were also close to 100%.

Intranasally delivered mAbs can potentially treat or prevent several diseases located in the brain or the respiratory tract, and the target region of the mAbs in the nasal cavity depends on the sites of the diseases and the transport mechanism of the mAbs. In the case of treating a disease located in the brain, pre-clinical studies in animals showed that intranasal delivery of antibodies or antibody-drug conjugates targeting the brain has the potential to treat glioblastoma or Alzheimer’s disease. Chu et al. functionalized temozolomide-loaded poly(lactic-glycolic acid) (PLGA) nanoparticles with anti-EPHA3 antibodies for nose-to-brain delivery (Chu et al., 2018). They demonstrated that intranasal administration could more efficiently deliver the drug into the brain of glioma-bearing rats than intravenous administration. Musumeci et al. utilized chitosan-coated PLGA nanoparticles or nanostructured lipid carriers (NLC) to carry mAb-neutralizing tumor necrosis factor-related apoptosis-inducing ligand (TRAIL) for the treatment of Alzheimer disease (Musumeci et al., 2022). Their results indicated that through intranasal administration, the drug complexes could efficiently reach the brain of the mice. Although these studies did not specify the targeted region in the nasal cavity for mAb deposition, a literature reporting the general transportation mechanisms of the pharmaceuticals from the nasal cavity to the brain is available. Patel et al. summarized that drugs can be transported *via* the olfactory, trigeminal, and systemic pathways (Patel, Patel and Wairkar, 2022). Specifically, olfactory epithelium should be targeted in the olfactory pathway, maxillary and ophthalmic nerves should be targeted in the trigeminal pathway, and the capillary blood vessels in the nasal mucosa should be targeted for the systemic pathway (Patel, Patel and Wairkar, 2022). It is likely that the deposition of the mAb powder at the olfactory region can be increased, if needed, by adjusting the insertion angle of the UDS Powder device.

In the case of respiratory tract infections, intranasal delivery of the mAb can passively immunize the respiratory tract mucosal surface and prevent viral infections (Mazanec et al., 1992; Weltzin and Monath, 1999). Ye et al. delivered mAb specific to influenza virus H5 hemagglutinin intranasally to the mice before or after they were inoculated with a sublethal dose of H5N1 influenza viruses. Their results indicated that intranasal delivery of mAb could provide protection against H5N1 influenza virus infections (Ye et al., 2010). Weltzin et al. delivered a mAb that neutralizes RSV intranasally to the mice. Their results indicated that intranasal delivery of mAb prior to RSV challenge can protect the mice from both nasal and lung infections. Interestingly, the protection against lung infection can last as long as 3 days, and the mAb was still present in the lungs for 4 days, while the protection against nasal infection of RSV cannot last for one day, which is likely because of the rapid clearance of the nasal mucus into the nasopharynx (Weltzin et al., 1994). In another study, Weltzin et al. delivered the same mAb intranasally to the Rhesus monkeys daily 2 days before and 4 days after RSV challenge and showed that the mAb protected the monkeys from infections (Weltzin et al., 1996). They also found that the mAb remained in nasal secretions for a few days after intranasal administration, while neutralizing concentration remained for more than one day after the treatment (Weltzin et al., 1996).

Recently, the outbreak of COVID-19 raised people’s interest in passive immunization by applying intranasal mAbs. Halwe et al. indicated that intranasal delivery of DZIF-10c, a neutralizing antibody against SARS-COV-2, can protect hACE2-transduced mice from infection (Halwe et al., 2021). However, they did not analyze the exact biodistribution of the DZIF-10c following intranasal administration. Ku et al. delivered an engineered immunoglobulin M (IgM) neutralizing antibody (IgM-14) to the nasal cavities of mice for the prophylactic treatment of COVID-19. To study the biodistribution of IgM-14 following intranasal administration, they labeled the IgM-14 with Alexa Fluor 750 (AF750) dye and applied an IVIS imager to determine its distribution *in vivo* and *ex vivo* (Ku et al., 2021). Their results show that following intranasal administration, IgM-14 has long-term retention in both the nasal cavity (168 h) and lung (96 h), which conferred protection against respiratory infection since nasal epithelium is the primary location where SARS-CoV-2 starts to infect, and the virus can be aspirated into the lungs (Hou et al., 2020; Ku et al., 2021).

In conclusion, all the mAb formulations in those mentioned examples are in liquid form, and the exploration of delivering mAb in powder form intranasally is still in its infancy. However, there is an example of intranasally delivering an antiviral agent in powder form. Seow et al. utilized spray drying and spray freeze-drying technology to produce powder formulations containing tamibarotene, a broad-spectrum antiviral agent against SARS-CoV-2 (Seow et al., 2022). By physically mixing the particles with sizes above 10 μm and below 5 μm, both nasal and lung deposition can be achieved by a single route of intranasal administration. However, the regulatory pathway is not clear for products that target both nasal cavity and lungs. In summary, to neutralize viruses, the posterior nasal cavity, except the olfactory region, is likely the key region to deliver the mAbs to, while deposition in the nasopharynx region, trachea, and lung should also be beneficial.

## 4. Conclusions

It is concluded that thin-film freeze-drying is suitable for converting mAbs into aerosolizable dry powders for intranasal delivery into the posterior nasal cavity using a dry powder spraying device such as Aptar’s UDS Powder device. With proper formulation compositions, subjecting mAbs such as AUG-3387 to thin-film freeze-drying did not cause an increase in high molecular weight species and the mass percentage of proteinaceous particles remained below 2% *w/w*. For intranasal liquid products, droplet/particle size distribution is considered a critical attribute. However, to measure the particle size distribution of an intranasal dry powder product, the traditional laser diffraction method used for intranasal liquid products may need to be modified to account for the challenge of identifying a stable phase and the low transmission of the sprayed powders.

## ACKNOWLEDGEMENTS

Z Cui and RO Williams III report financial support from TFF Pharmaceuticals, Inc. The UDS Powder device was generously provided by the AptarGroup, Inc. YS Yu was supported in part by the Y.L. Lin Hung Tai Education Foundation, and YS Yu and KCW Wu report support from the Taiwan Ministry of Science and Technology (award #: 111- 2628-E-002-008). K AboulFotouh was supported in part by an Egyptian Government Fellowship (GM 1105). We thank Dr. Hugh Smyth in the College of Pharmacy at UT Austin for the laser sheet system used to acquire the spray pattern and plume geometry data. The graphical abstract was created with BioRender.com and adopted from “Head and Neck Anatomy”, by BioRender.com (2020). Retrieved fromhttps://app.biorender.com/biorender-templates

## CRediT Author Statement

YS Yu: Conceptualization, Methodology, Investigation, Visualization, Formal analysis, Writing Original Draft; K AboulFotouh: Methodology, investigation, Review & Editing; H Xu, S Sahakijpijarn: Methodology, investigation; G Williams, J Suman, C Cano: Conceptualization, Resources, Writing-Review & Editing; Z Warnken: Methodology, investigation, Writing-Review & Editing; KCW Wu: Writing-Review & Editing, Funding acquisition; RO Williams III: Resources, Writing-Review & Editing, Funding acquisition; Z Cui: Conceptualization, Resources, Validation, Writing-Review & Editing, Supervision, Funding acquisition.

## References

1. Allison, S.D., Molina, M.d.C., Anchordoquy, T.J., 2000. Stabilization of lipid/DNA complexes during the freezing step of the lyophilization process: the particle isolation hypothesis. Biochim. Biophys. Acta Biomembr. 1468, 127–138. 10.1016/S0005-2736(00)00251-0.

2. Barnard, J.G., Singh, S., Randolph, T.W., Carpenter, J.F., 2011. Subvisible particle counting provides a sensitive method of detecting and quantifying aggregation of monoclonal antibody caused by freeze-thawing: Insights into the roles of particles in the protein aggregation pathway. J. Pharm. Sci. 100, 492–503. 10.1002/jps.22305.

3. Chu, L., Wang, A., Ni, L., Yan, X., Song, Y., Zhao, M., Sun, K., Mu, H., Liu, S., Wu, Z., Zhang, C., 2018. Nose-to-brain delivery of temozolomide-loaded PLGA nanoparticles functionalized with anti-EPHA3 for glioblastoma targeting. Drug. Deliv. 25, 1634–1641. 10.1080/10717544.2018.1494226.

4. Cleland, J.L., Lam, X., Kendrick, B., Yang, J., Yang, T.H., Overcashier, D., Brooks, D., Hsu, C., Carpenter, J.F., 2001. A specific molar ratio of stabilizer to protein is required for storage stability of a lyophilized monoclonal antibody. J. Pharm. Sci. 90, 310–321. 10.1002/1520-6017(200103)90:3<310::aid-jps6>3.0.co;2-r.

5. Coll, M., Solans, X., Font-Altaba, M., Subirana, J., 1986. Structure of L-leucine: a redetermination. Acta Crystallogr. C 42, 599–601. 10.1107/S0108270186095240.

6. Djupesland, P.G., 2013. Nasal drug delivery devices: characteristics and performance in a clinical perspective-a review. Drug. Deliv. Transl. Res. 3, 42–62. 10.1007/s13346-012-0108-9.

7. Emami, F., Vatanara, A., Park, E.J., Na, D.H., 2018. Drying technologies for the stability and bioavailability of biopharmaceuticals. Pharmaceutics 10. 10.3390/pharmaceutics10030131.

8. Emig, C.J., Mena, M.A., Henry, S.J., Vitug, A., Ventura, C.J., Fox, D., Nguyenla, X.H., Xu, H., Moon, C., Sahakijjpijarn, S., Kuehl, P.J., Revelli, D., Cui, Z., Williams, R.O., Christensen, D.J., 2021. AUG-3387, a human-derived monoclonal antibody neutralizes SARS-CoV-2 variants and reduces viral load from therapeutic treatment of hamsters in vivo. bioRxiv, 2021.2010.2012.464150. 10.1101/2021.10.12.464150.

9. Engstrom, J.D., Lai, E.S., Ludher, B.S., Chen, B., Milner, T.E., Williams, R.O., Kitto, G.B., Johnston, K.P., 2008. Formation of stable submicron protein particles by thin film freezing. Pharm. Res. 25, 1334–1346. 10.1007/s11095-008-9540-4.

10. Filipović-Grčić, J., Hafner, A., Nasal powder drug delivery, Pharm. Sci. Encyclopedia, pp. 1–32. 10.1002/9780470571224.pse356.

11. Fronczek, F.R., Kamel, H.N., Slattery, M., 2003. Three polymorphs (alpha, beta, and delta) of D-mannitol at 100 K. Acta Crystallogr. C 59, O567–570. 10.1107/s0108270103018961.

12. Glücklich, N., Dwivedi, M., Carle, S., Buske, J., Mäder, K., Garidel, P., 2020. An in-depth examination of fatty acid solubility limits in biotherapeutic protein formulations containing polysorbate 20 and polysorbate 80. Int. J. Pharm. 591, 119934. 10.1016/j.ijpharm.2020.119934.

13. Guo, S., Yu, C., Guo, X., Jia, Z., Yu, X., Yang, Y., Guo, L., Wang, L., 2022. Subvisible particle analysis of 17 monoclonal antibodies approved in China using flow imaging and light obscuration. J. Pharm. Sci. 111, 1164–1171. 10.1016/j.xphs.2021.09.021.

14. Haeuser, C., Goldbach, P., Huwyler, J., Friess, W., Allmendinger, A., 2020. Excipients for room temperature stable freeze-dried monoclonal antibody formulations. J. Pharm. Sci. 109, 807–817. 10.1016/j.xphs.2019.10.016.

15. Halwe, S., Kupke, A., Vanshylla, K., Liberta, F., Gruell, H., Zehner, M., Rohde, C., Krähling, V., Gellhorn Serra, M., Kreer, C., Klüver, M., Sauerhering, L., Schmidt, J., Cai, Z., Han, F., Young, D., Yang, G., Widera, M., Koch, M., Werner, A., Kämper, L., Becker, N., Marlow, M.S., Eickmann, M., Ciesek, S., Schiele, F., Klein, F., Becker, S., 2021. Intranasal administration of a monoclonal neutralizing antibody protects mice against SARS-CoV-2 infection. Viruses 13, 1498. 10.3390/v13081498.

15a. Heo, Y.-A., 2022. Sotrovimab: First approval. Drugs 82, 477-484. 10.1007/s40265-022-01690-7.

16. Hou, Y.J., Okuda, K., Edwards, C.E., Martinez, D.R., Asakura, T., Dinnon, K.H., 3rd, Kato, T., Lee, R.E., Yount, B.L., Mascenik, T.M., Chen, G., Olivier, K.N., Ghio, A., Tse, L.V., Leist, S.R., Gralinski, L.E., Schäfer, A., Dang, H., Gilmore, R., Nakano, S., Sun, L., Fulcher, M.L., Livraghi-Butrico, A., Nicely, N.I., Cameron, M., Cameron, C., Kelvin, D.J., de Silva, A., Margolis, D.M., Markmann, A., Bartelt, L., Zumwalt, R., Martinez, F.J., Salvatore, S.P., Borczuk, A., Tata, P.R., Sontake, V., Kimple, A., Jaspers, I., O’Neal, W.K., Randell, S.H., Boucher, R.C., Baric, R.S., 2020. SARS-CoV-2 reverse genetics reveals a variable infection gradient in the respiratory tract. Cell 182, 429-446.e414. 10.1016/j.cell.2020.05.042.

17. Hufnagel, S., Xu, H., Sahakijpijarn, S., Moon, C., Chow, L.Q.M., Williams Iii, R.O., Cui, Z., 2022. Dry powders for inhalation containing monoclonal antibodies made by thin-film freeze-drying. Int. J. Pharm. 618, 121637. 10.1016/j.ijpharm.2022.121637.

18. Jain, H., Schweitzer, J.W., Justice, N.A., 2022. Respiratory syncytial virus infection, StatPearls. StatPearls Publishing, Treasure Island (FL).

19. Kalonia, C., Kumru, O.S., Indira Prajapati, V., Mathaes, R., Engert, J., Zhou, S., Russell Middaugh, C., Volkin, D.B., 2015. Calculating the mass of subvisible protein particles with improved accuracy using microflow imaging data. J. Pharm. Sci. 104, 536–547. 10.1002/jps.24156.

20. Keam, S.J., 2022. Tixagevimab + cilgavimab: First approval. Drugs 82, 1001–1010. 10.1007/s40265-022-01731-1.

21. Kelley, B., De Moor, P., Douglas, K., Renshaw, T., Traviglia, S., 2022. Monoclonal antibody therapies for COVID-19: Lessons learned and implications for the development of future products. Curr. Opin. Biotechnol. 78, 102798. 10.1016/j.copbio.2022.102798.

22. Ku, Z., Xie, X., Hinton, P.R., Liu, X., Ye, X., Muruato, A.E., Ng, D.C., Biswas, S., Zou, J., Liu, Y., Pandya, D., Menachery, V.D., Rahman, S., Cao, Y.A., Deng, H., Xiong, W., Carlin, K.B., Liu, J., Su, H., Haanes, E.J., Keyt, B.A., Zhang, N., Carroll, S.F., Shi, P.Y., An, Z., 2021. Nasal delivery of an IgM offers broad protection from SARS-CoV-2 variants. Nature 595, 718–723. 10.1038/s41586-021-03673-2.

23. Li, L., Gorukanti, S., Choi, Y.M., Kim, K.H., 2000. Rapid-onset intranasal delivery of anticonvulsants: pharmacokinetic and pharmacodynamic evaluation in rabbits. Int. J. Pharm. 199, 65–76. 10.1016/S0378-5173(00)00373-2.

24. Madden, J., McGandy, E., Seeman, N., Harding, M., Hoy, A., 1972. The crystal structure of the monoclinic form of L-histidine. Acta Crystallogr. B 28, 2382–2389. 10.1107/S056774087200617X.

25. Madden, J.J., McGandy, E.L., Seeman, N.C., 1972. The crystal structure of the orthorhombic form of L-(+)-histidine. Acta Crystallogr. B 28, 2377–2382. 10.1107/S0567740872006168.

26. Malvern Panalytical, 2005. Nasal Spray Device Measurement Using Laser Diffraction and the Spraytec. AZO MATERIALS.

27. Mazanec, M.B., Lamm, M.E., Lyn, D., Portner, A., Nedrud, J.G., 1992. Comparison of IgA versus IgG monoclonal antibodies for passive immunization of the murine respiratory tract. Virus Res. 23, 1–12. 10.1016/0168-1702(92)90063-f.

28. Mohammed, A.R., Coombes, A.G.A., Perrie, Y., 2007. Amino acids as cryoprotectants for liposomal delivery systems. Eur. J. Pharm. Sci. 30, 406–413. 10.1016/j.ejps.2007.01.001.

29. Moreira, T.G., Matos, K.T.F., De Paula, G.S., Santana, T.M.M., Da Mata, R.G., Pansera, F.C., Cortina, A.S., Spinola, M.G., Baecher-Allan, C.M., Keppeke, G.D., Jacob, J., Palejwala, V., Chen, K., Izzy, S., Healey, B.C., Rezende, R.M., Dedivitis, R.A., Shailubhai, K., Weiner, H.L., 2021. Nasal administration of anti-CD3 monoclonal antibody (foralumab) reduces lung inflammation and blood inflammatory biomarkers in mild to moderate COVID-19 patients: A pilot study. Front. Immunol. 12, 709861. 10.3389/fimmu.2021.709861.

30. Musumeci, T., Di Benedetto, G., Carbone, C., Bonaccorso, A., Amato, G., Lo Faro, M.J., Burgaletto, C., Puglisi, G., Bernardini, R., Cantarella, G., 2022. Intranasal administration of a TRAIL neutralizing monoclonal antibody adsorbed in PLGA nanoparticles and NLC nanosystems: An in vivo study on a mouse model of Alzheimer’s disease. Biomedicines 10. 10.3390/biomedicines10050985.

31. Nižić Nodilo, L., Ugrina, I., Špoljarić, D., Amidžić Klarić, D., Jakobušić Brala, C., Perkušić, M., Pepić, I., Lovrić, J., Saršon, V., Safundžić Kučuk, M., 2021. A dry powder platform for nose-to-brain delivery of dexamethasone: Formulation development and nasal deposition studies. Pharmaceutics 13, 795. 10.3390/pharmaceutics13060795.

32. Pabari, R.M., Ryan, B., McCarthy, C., Ramtoola, Z., 2011. Effect of microencapsulation shear stress on the structural integrity and biological activity of a model monoclonal antibody, trastuzumab. Pharmaceutics 3, 510–524. 10.3390/pharmaceutics3030510.

33. Patel, D., Patel, B., Wairkar, S., 2022. Intranasal delivery of biotechnology-based therapeutics. Drug Discov. Today 27, 103371. 10.1016/j.drudis.2022.103371.

34. Petersen, E., Koopmans, M., Go, U., Hamer, D.H., Petrosillo, N., Castelli, F., Storgaard, M., Al Khalili, S., Simonsen, L., 2020. Comparing SARS-CoV-2 with SARS-CoV and influenza pandemics. Lancet Infect. Dis. 20, e238–e244. 10.1016/s1473-3099(20)30484-9.

35. Pormohammad, A., Ghorbani, S., Khatami, A., Razizadeh, M.H., Alborzi, E., Zarei, M., Idrovo, J.-P., Turner, R.J., 2021. Comparison of influenza type A and B with COVID- 19: A global systematic review and meta-analysis on clinical, laboratory and radiographic findings. Rev. Med. Virol. 31, e2179. 10.1002/rmv.2179.

36. Praphawatvet, T., Cui, Z., Williams, R.O., 2022. Pharmaceutical dry powders of small molecules prepared by thin-film freezing and their applications – A focus on the physical and aerosol properties of the powders. Int. J. Pharm. 629, 122357. 10.1016/j.ijpharm.2022.122357.

37. Sahakijpijarn, S., Moon, C., Koleng, J.J., Christensen, D.J., Williams Iii, R.O., 2020. Development of remdesivir as a dry powder for inhalation by thin film freezing. Pharmaceutics 12. 10.3390/pharmaceutics12111002.

38. Seow, H.C., Liao, Q., Lau, A.T.Y., Leung, S.W.S., Yuan, S., Lam, J.K.W., 2022. Dual targeting powder formulation of antiviral agent for customizable nasal and lung deposition profile through single intranasal administration. Int. J. Pharm. 619, 121704. 10.1016/j.ijpharm.2022.121704.

39. Sharma, D.K., King, D., Oma, P., Merchant, C., 2010a. Micro-flow imaging: flow microscopy applied to sub-visible particulate analysis in protein formulations. AAPS J. 12, 455–464. 10.1208/s12248-010-9205-1.

40. Sharma, D.K., Oma, P., Pollo, M.J., Sukumar, M., 2010b. Quantification and characterization of subvisible proteinaceous particles in opalescent mAb formulations using micro-flow imaging. J. Pharm. Sci. 99, 2628–2642. 10.1002/jps.22046.

41. US FDA, 2002. Nasal spray and inhalation solution, suspension, and spray drug products — Chemistry, manufacturing, and controls documentation.

42. US FDA, 2003. Bioavailability and bioequivalence studies for nasal aerosols and nasal sprays for local action.

43. USP-NF, 2023. 〈787〉 Subvisible Particulate Matter in Therapeutic Protein Injections, United States Pharmacopeia, Rockville, MD. 10.31003/USPNF_M6497_02_01.

44. van Riel, D., den Bakker, M.A., Leijten, L.M.E., Chutinimitkul, S., Munster, V.J., de Wit, E., Rimmelzwaan, G.F., Fouchier, R.A.M., Osterhaus, A.D.M.E., Kuiken, T., 2010. Seasonal and pandemic human influenza viruses attach better to human upper respiratory tract epithelium than avian influenza viruses. Am. J. Clin. Pathol. 176, 1614–1618. 10.2353/ajpath.2010.090949.

45. Wang, J.-L., Kuang, M., Xu, H., Williams, R.O., Cui, Z., 2023. Accelerated water removal from frozen thin films containing bacteria. Int. J. Pharm. 630, 122408. 10.1016/j.ijpharm.2022.122408.

46. Warnken, Z.N., Smyth, H.D.C., Davis, D.A., Weitman, S., Kuhn, J.G., Williams, R.O., III, 2018. Personalized medicine in nasal delivery: The use of patient-specific administration parameters to improve nasal drug targeting using 3D-printed nasal replica casts. Mol. Pharm. 15, 1392–1402. 10.1021/acs.molpharmaceut.7b00702.

47. Weltzin, R., Hsu, S.A., Mittler, E.S., Georgakopoulos, K., Monath, T.P., 1994. Intranasal monoclonal immunoglobulin A against respiratory syncytial virus protects against upper and lower respiratory tract infections in mice. Antimicrob. Agents Chemother. 38, 2785–2791. 10.1128/aac.38.12.2785.

48. Weltzin, R., Monath, T.P., 1999. Intranasal antibody prophylaxis for protection against viral disease. Clin. Microbiol. Rev. 12, 383–393. 10.1128/cmr.12.3.383.

49. Weltzin, R., Traina-Dorge, V., Soike, K., Zhang, J.Y., Mack, P., Soman, G., Drabik, G., Monath, T.P., 1996. Intranasal monoclonal IgA antibody to respiratory syncytial virus protects rhesus monkeys against upper and lower respiratory tract infection. J. Infect. Dis. 174, 256–261. 10.1093/infdis/174.2.256.

50. Ye, J., Shao, H., Hickman, D., Angel, M., Xu, K., Cai, Y., Song, H., Fouchier, R.A., Qin, A., Perez, D.R., 2010. Intranasal delivery of an IgA monoclonal antibody effective against sublethal H5N1 influenza virus infection in mice. Clin. Vaccine Immunol. 17, 1363–1370. 10.1128/cvi.00002-10.

51. Yu, Y.S., AboulFotouh, K., Xu, H., Williams, G., Suman, J., Cano, C., Warnken, Z.N., Wu, K.C., Williams, R.O., 3rd, Cui, Z., 2023. Feasibility of intranasal delivery of thin-film freeze-dried, mucoadhesive vaccine powders. Int. J. Pharm. 640, 122990. 10.1016/j.ijpharm.2023.122990.

52. Zhang, Y., Soto, M., Ghosh, D., Williams, R.O., 2021. Manufacturing Stable Bacteriophage Powders by Including Buffer System in Formulations and Using Thin Film Freeze-drying Technology. Pharm. Res. 38, 1793–1804. 10.1007/s11095-021-03111-y.

